# Human neutrophil development and functionality are enabled in a humanized mouse model

**DOI:** 10.1101/2022.02.25.481352

**Authors:** Yunjiang Zheng, Esen Sefik, John Astle, Angel G. Solis, Hasan H. Öz, Ruaidhrí Jackson, Emanuela M. Bruscia, Stephanie Halene, Liang Shan, Richard A Flavell

## Abstract

Mice with a functional human immune system serve as an invaluable tool to study the development and function of human immune system in vivo. A major technological limitation of all current humanized mouse models is the lack of mature and functional human neutrophils in circulation and tissues. To overcome this, we generated a humanized mouse model named MISTRGGR, in which the mouse granulocyte colony stimulating factor (G-CSF) was replaced with human G-CSF and the mouse G-CSF receptor gene was deleted (G-CSFR) in existing MISTRG mice. By targeting the GCSF cytokine-receptor axis, we dramatically improved the reconstitution of mature circulating and tissue-infiltrating human neutrophils in MISTRGGR mice. Moreover, these functional human neutrophils in MISTRGGR are recruited upon inflammatory and infectious challenges and help reduce bacterial burden. MISTRGGR mice represent a unique mouse model that finally permits the study of human neutrophils in health and disease.

**Key Points:**

- Targeting the GCSF cytokine-receptor axis dramatically improves circulating and tissue-infiltrating human neutrophils in MISTRGGR mice.
- Human neutrophils generated in MISTRGGR mice are functional and are able to respond robustly to inflammatory and infectious stimuli.

## Introduction

Mice serve as valuable tools for the study of *in vivo* immune responses. Although the overall structure and composition of the murine and human immune systems are comparable at many levels, extensive evolutionary pressures over millions of years have resulted in significant changes^1-4^. Differences between the human and murine immune systems have been widely characterized in blood. Transcriptional analysis of mouse and human cells show similarities in lineage defining genes and aspects of cell identity, yet many human and mouse immune cells show striking differences in their activities and effector or regulatory mechanisms^5^. Neutrophils are one such cell type for which human or murine specific effects drive differences in immunity and pathogenesis.

Neutrophils are indispensable during injury, infection and as more recently appreciated in homeostatic immune surveillance in both humans and mice^6-9^. They are the most abundant circulating leukocytes and play essential roles in fending against microbial infections through a variety of mechanism including phagocytosis, release of reactive oxygen species (ROS), antimicrobial peptides and proteases, and by their unique formation of neutrophil extracellular traps (NETs)^9,10^. Both human and mouse neutrophils produce various cytokines and chemokines to influence inflammation and regulate adaptive immune responses^11^. Yet, there are significant differences in the maturation, migration and function of human and mouse neutrophils. The structure of key neutrophil molecules (selectins, FcαRI, serine proteases)^12-15^, expression of cytokines (IL10, IL17)^16,17^, activation pathways of ROS^18,19^ and cytotoxic granular content (defensins, bactericidal enzymes)^14,20-22^ display major cross-species alterations that can affect migration, signaling, effector mechanisms and neutrophil activation^23^. These human to mouse differences in the overall biology of neutrophils have striking implications for immunity against infections, pathogenesis of cancer, and health^4,6-23^.

Neutrophils are pivotal to the human innate defense against various bacterial and fungal pathogens, including *Listeria monocytogenes, Pseudomonas aeruginosa* and *Candida auris*^*24-27*^. In contrast to mice, for which neutrophil deficiencies do not cause a progressive health deterioration under specific-pathogen-free (SPF) conditions^28^, neutropenic humans suffer greatly from exposure to microbials in daily life^29^, suggesting an even more crucial role of neutrophils in immune defense against infections in humans. Individuals with severe congenital neutropenia, as well as patients undergoing chemotherapy and bone marrow transplantation, are commonly treated with recombinant human G-CSF (filgrastim). G-CSF drives granulopoiesis and improves and accelerates the recovery of circulating neutrophil counts, thus reducing the frequency and severity of infections^30,31^.

In various human and murine tumors, neutrophils compose a significant part of tumor-infiltrating myeloid cells and can display both pro and anti-tumor effects^23,32,33^. Although recent work has provided clues to how neutrophils play a complex role in pathologies such as cancer progression, significant gaps remain in understanding the regulation of neutrophil infiltration and survival within the tumor microenvironment and the switch between pro- and anti-tumor phenotypes.

Neutrophils can infiltrate multiple tissues like the spleen and lung even at the steady-state and participate in local immune homeostasis^6-8,34^. Yet, it remains unknown to what extent infiltrating neutrophils contribute to homeostatic immune surveillance and tissue physiology, especially in humans, as approaches to study and manipulate human neutrophils in human tissues are limited. Conflicting reports in mouse studies and a general lack of data about the function of human neutrophils have created a fundamental need for development of humanized mice that allow for the development and maturation of functional human neutrophils, that are capable of response to inflammation.

Development of humanized hemato-lymphoid system mice (referred here as humanized mice) that are generated by transplantation of human hematopoietic cells into various strains of immune-compromised mice have provided a leap forward for studying human immune function in vivo. MISTRG mice, which harbor human cytokines MCSF, IL3/GMCSF, THPO and human SIRPα have highly improved maturation and function of human monocytes, macrophages, NK cells as well as the general humanization of the immune system^35^. Strikingly however, no system that supports circulating human neutrophils currently exists. Here, we describe a novel humanized mouse model by humanization of the cytokine G-CSF and ablation of the murine G-CSF receptor (G-CSFR), to finally allow human neutrophil development and survival in an in vivo model system. With successful human neutrophil reconstitution in the periphery, this humanized mouse model can now be applied to study human neutrophils and unravel their major contribution to health and disease.

## Materials and Methods

### Transplantation of human CD34+ hematopoietic progenitor cells into mice

Fetal liver samples were cut in small fragments, treated for 45 min at 37 °C with collagenase D (Roche, 200 μg/ml), and prepared into a cell suspension. Human CD34+ cells were purified by performing density gradient centrifugation (Lymphocyte Separation Medium, MP Biomedicals), followed by positive immunomagnetic selection with EasySep™ Human CD34 Positive Selection Kit (Stemcell). For intra-hepatic engraftment, newborn 1-3 day-old pups were sublethally irradiated at a dose of 150 Rads (X-RAD 320 irradiator), and unless otherwise specified, 20,000 fetal liver CD34+ cells in 20 μl of PBS were injected into the liver with a 22-gauge needle (Hamilton Company). All use of human materials was approved by the Yale University Human Investigation Committee.

### Flow Cytometry

All mice were analyzed at approximately 6-8 weeks post-engraftment. Blood was collected retro-orbitally. Single cell suspensions were prepared from bone marrow and lung. Samples were stained with fluorochrome-labeled monoclonal antibodies against mouse and human cell-surface antigens and analyzed on an LSRII flow cytometer (BDBiosciences). Data were analyzed using FloJo software.

### *In vitro* phagocytosis assay

Protocols as previously described^36^ were adapted and optimized. Blood was incubated with pHrodo™ Green *E. coli* BioParticles® Conjugate (Life Technologies, 1mg/ml) in a 96-well plate at 37 °C for 0, 20 and 90 minutes respectively (10 ug *E*.*coli* particles per 50ul blood). Cells were fixed at an added final concentration of 1.6% PFA and stained for lineage characterization. All buffers were tested for pH 7.4 prior to use.

### LPS nebulization, in vivo labelling, and tissue collection

Mice were exposed to aerosolized lipopolysaccharide (LPS from *P. aeruginosa* 10, Sigma L9143) using a nebulizer (Pulmo-Aide compressor) as previously described^37^. LPS (12.5 mg) dissolved in 5.0 ml of PBS were administered using a nebulizer connected to a container with vent holes over 15 min. At 24 hours after nebulization, mice were anesthetized. To distinguish between circulating and lung resident or infiltrating human immune cells, fluorochrome labeled anti-human CD45 antibody (2.5μg, HI30) was intravenously injected 5 min prior to sample collection. BAL was performed using standard methods with a 22G Catheter (BD)^38^. Mice were then euthanized for tissue collection as previously described.

### *Pseudomonas aeruginosa* infection

Two doses of 100 ug anti-Ly6G antibody (clone: 1A8, BioXCell) in 100 ul PBS were intravenously injected to recipient mice at approximately 48 hours and 3 hours prior to the intranasal admiration of *Pseudomonas aeruginosa*. Prior to inoculation (∼24hr), *P. aeruginosa* (PA14) were grown at 37□°C in a shaking incubator for approximately 16 hr in LB medium. Bacteria were subcultured in a 1:20 ratio until OD600 was between 1.5 and 2.0, which is approximately 1.5 × 10^9^ and 2 × 10^9^ CFU/ml. Mice were anesthetized with methoxyflurane, and a dose of 500 or 5000 CFU in 20 ul PBS was instilled intranasally using a micropipette. Mice were sacrificed at 18 or 24 hours after infection, unless otherwise stated. Livers and lungs were collected sterilely, homogenized with gentleMACS Dissociator (Miltenyi), plated on LB plates and cultured at 37□°C overnight for the calculation of bacterial loads. Alternately, a subset of mice were used to characterize of the immune cells by flow cytometry as previously described.

### Statistical analysis

Statistical analysis was performed with the GraphPad Prism 7 software, using two-tailed unpaired Student’s t-test (Mann-Whitney test).

## Results

Granulocyte colony-stimulating factor (G-CSF) is the major cytokine that regulates the commitment, development, maturation, and survival of neutrophilic granulocytes^39-42^. Deletion of the murine *G-CSF* gene revealed physiologic roles for this cytokine in granulopoiesis at both steady state and condition of inflammation and infections^40^. G-CSF is produced by a variety of cells including macrophages, T cells, endothelial cells, and fibroblasts upon inflammation signals, and in turn induces proliferation of neutrophilic progenitors and the release of post-mitotic neutrophils from the bone marrow^43,44^. Human and murine G-CSF proteins display a relatively high degree of amino acid sequence homology (73%)^45^. Early studies have shown receptor-binding cross-reactivity between human and mouse G-CSF^46^; however, little is known of the efficacy of these proteins in a cross-species bone marrow microenvironment.

Given the crucial role of G-CSF in regulating granulopoiesis, we hypothesized that humanization of cytokine G-CSF will favor human neutrophil reconstitution in humanized MISTRG mice. Thus, we generated mice that are deficient in mouse *G-CSF (Csf3)* and express human *GCSF (CSF3)* instead using CRISPR-Cas9 system^47^, referred to as MISTRGG (G for G-CSF^hh^) mice hereafter. The construct was designed to replace the entire mouse *G-CSF* open reading frame (ORF) (∼1.6kb) with the human ORF, but to maintain the promotor, 5’ and 3’ UTRs primarily of mouse origin (Fig. 1a). We confirmed the loss of mouse G-CSF and expression of the human G-CSF in the bone marrow and blood plasma after LPS stimulation (Fig. 1b, c), a widely-accepted protocol that induces systemic G-CSF expression^48^. To assess the effect of G-CSF humanization on granulocyte reconstitution in humanized mice, newborn MISTRGG and control MISTRG mice were intra-hepatically injected with human fetal CD34+ hematopoietic stem and progenitor cells. As a readout of engraftment, human and mouse immune cells were quantified at 7-8 weeks post-injection (Fig. S1). Replacement of the mouse *G-CSF* with the human *GCSF* yielded increased human neutrophil frequencies in the bone marrow at steady state but did not impact circulating neutrophils in blood (Fig. 1d). Other major immune human populations such as T, NK, B and CD33^+^ myeloid cells remained unaffected in MISTRGG mice (Fig. 1e). The levels of murine neutrophils in circulation and bone marrow were not affected (Fig. 1d), which we hypothesize is a result of the functionality of human G-CSF on the mouse G-CSFR due to the reported capability of recombinant human G-CSF to induce murine granulocytic colony formation^46,49^.

**Figure 1.**
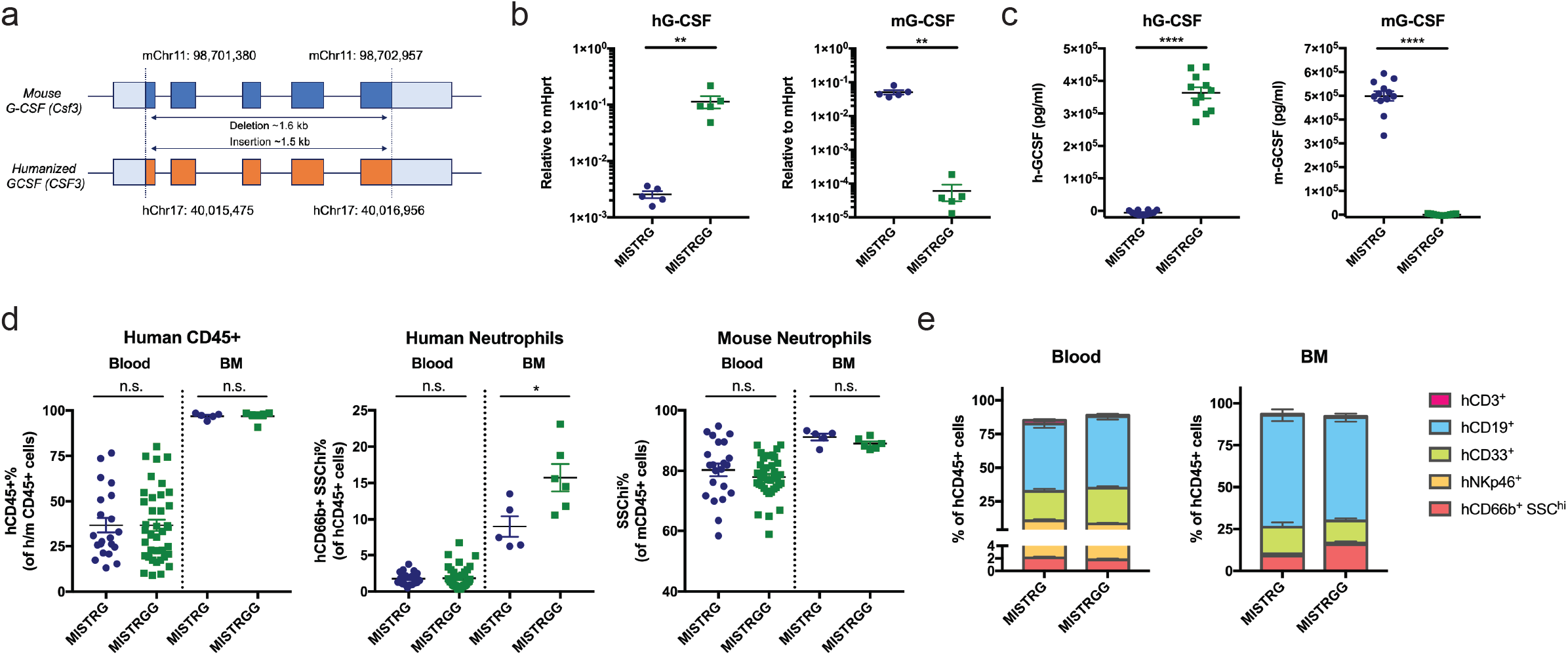
G-CSF humanization enhances neutrophil commitment in the bone marrow. (a) Schematic design of human GCSF knock-in replacement. The entire mouse open reading frame (in blue) is replaced by human open reading frame (in orange). Both 5’ and 3’ UTR sequences are primarily murine. (b) Relative expression of human and mouse G-CSF mRNA in the bone marrow at 3 hours (hr) after LPS administration (50 ug, i.p.). (c) Human and mouse G-CSF proteins measured in plasma at 3hr after LPS administration (50 ug, i.p.). (d) Frequencies of human hematopoietic cells (hCD45+), human neutrophils (hCD66b+ SSChi), and mouse neutrophils (mCD45+ SSChi) in the blood and bone marrow. (e) Frequencies of human lineages in the blood (MISTRG, n=14; MISTRGG, n=20) and the bone marrow (MISTRG, n=5; MISTRGG, n=6) of mice at 7 weeks post-engraftment. Data pooled from at least 3 independent experiments. Mice were irradiated with 150 Rads and intra-hepatically injected with 20,000 human CD34+ cells at 1-3 days after birth.

Therefore, in order to reduce competition between human and murine granulocytic progenitors over niche space, human G-CSF and other cytokines and growth factors, we deleted mouse G-CSF receptor (*G-CSFR*, also known as *Csf3r/Cd114*) in MISTRGG mice, referred to as MISTRGGR (R for *G-CSF* R^-/-^) mice hereafter, as G-CSFR deficiency is expected to cause an 80% decrease in mouse neutrophils in circulation as well as the bone marrow^50^. We designed CRISPR guides targeting both the start codon and in the beginning of 3’ UTR, resulting in a deletion of ∼16.8kb spanning the entire mouse ORF (Fig. 2a). The loss of G-CSFR was confirmed at transcript level (Fig. 2b). To assess the outcome of G-CSFR deletion on mouse neutrophils, we quantified mouse neutrophils by flow cytometry in MISTRGGR^-/-^ mice. As anticipated, MISTRGGR^-/-^ mice had a significant reduction of mouse neutrophils in blood, bone marrow and peripheral tissues such as lungs (Fig. 2c).

**Figure 2.**
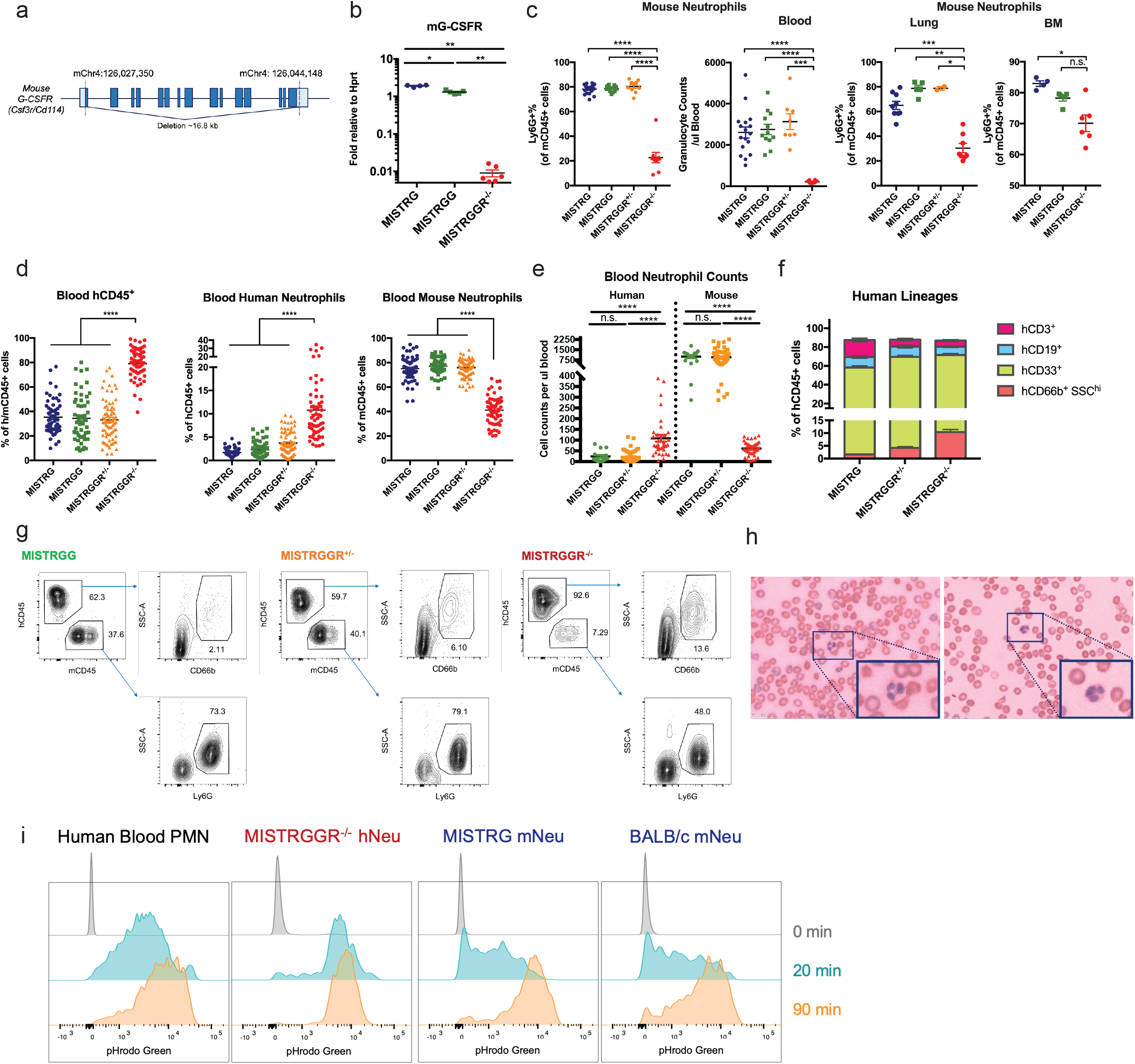
Depletion of murine neutrophils by G-CSFR ablation improves engraftment of human neutrophils in circulation. (a) Schematic design of mouse G-CSFR deletion. The entire G-CSFR open reading frame and first ∼300 bp of 3’ UTR are deleted. (b) Relative expression of G-CSFR in bone marrow of MISTRGGR-/-mice. Hprt was used as a housekeeping gene. (c) Frequencies and numbers of mouse granulocytes (Ly6G+) in blood (left), bone marrow (right) and lung (mid) of MISTRG, MISTRGG, MISTRGGR+/-and MISTRG-GR-/- mice. Whole lung tissues were analyzed without perfusion. (d - f) Characterization of human engraftment in the blood of MISTRG, MISTRGG, MISTRGGR+/-and MISTRGGR-/- mice at 7 weeks post-engraftment. Data pooled from at least 8 independent experiments. Mice were irradiated with 150 Rads and intra-hepatically injected with 15,000 ∼ 30,000 human fetal liver CD34+ cells at 1-3 days after birth. (d) Frequencies of human hematopoietic cells (hCD45+), human neutrophils (hCD66b+ SSChi), and mouse neutrophils (Ly6G+). (e) Numbers of human and mouse neutrophils. (f) Frequencies of human immune lineages (MISTRG, n=58; MISTRGGR+/-, n=38; MISTRGGR-/-, n=48). (g) Representative flow cytometry gating of human neutrophils in the blood of engrafted mice. (h) May-Grunwald Giemsa staining of engrafted MISTRGGR-/- blood smears. Neutrophils from two representative mice are shown in the enlarged box. (i) Representative flow cytometry analysis of phagocytosis by human blood PMNs (left), reconstituted human neutrophils from the blood of engrafted MISTRGGR-/- (middle left), mouse neutrophils from the blood of engrafted MISTRG (middle right) and unengrafted BALB/c (right). Blood samples were incubated with pHrodo™ Green E. coli BioParticles® Conjugate (10ug per 50ul blood) for the indicated times and fluorescent signals were analyzed by flow cytometry. Upper panel: 0 min. Middle panel: 20 min. Bottom panel: 90 min. Data were represenative of at least 2 independent experiments.

Given the reduced number of mouse neutrophils in MISTRGGR^-/-^, we wanted to test whether circulating human neutrophil proportions and numbers were affected in engrafted mice. Newborn MISTRGGR^-/-^ and control littermates were again injected with fetal CD34+ cells, and human engraftment were assessed similarly as in Fig. 1. Remarkably, overall humanization quantified by human CD45^+^ expression was enhanced in MISTRGGR^-/-^ compared with MISTRGGR^+/-^, MISTRGG and MISTRG groups. This enhanced humanization corresponded to a marked increase in human neutrophils (CD66b^+^ SSC^hi^) as well as an accompanied reduction in mouse neutrophils (Ly6G^+^) (Fig. 2d, e, g). The reconstitution of other major immune human populations such as B cells and CD33^+^ myeloid cells was unaffected (Fig. 2f, S2a). A definitive identification of human neutrophils was achieved by May-Grunwald Giemsa (MGG) staining in blood smears (Fig. 2h). These cells (as seen in the enlarged boxes) had segmented nuclei with approximately 3 lobes and a light pink cytoplasm, which is in accordance with the distinct morphology of mature human neutrophils^51^. We further tested the capacity of reconstituted human neutrophils to phagocytose inflammatory targets in an in vitro assay using fluorogenic *E. Coli* bioparticles^36^. Human neutrophils from MISTRGGR^-/-^ blood were capable of phagocytosis with efficacy similar to PMNs from human blood (Fig. 2i). These observations were specific to human and mouse neutrophils and were not observed in other lineages capable of phagocytosis such as human B cells (Fig. 2i, S2b).

One explanation for enhanced circulating neutrophils was enhanced neutrophil development in the bone marrow. While the overall human engraftment (hCD45^+^) in the bone marrow was comparable across different genotypes, MISTRGGR^-/-^ mice had significantly more human neutrophils (CD66b^+^ CD15^+^ SSC^hi^) (Fig. 3a, b, S3a). This population of neutrophils and neutrophilic progenitors have been shown to be further separated into CD49d^+^ 101^-^ pre-Neutrophils (Pre-Neu), primarily myelocytes and metamyelocytes, and CD101^+^ Neutrophils (Neu), consisting band and segmented neutrophils^52^. We’ve performed similar flow cytometric analysis and revealed that G-CSFR ablation improved reconstitution of both pre-Neu and Neu subsets (Fig. 3b, c), indicating an enhancement throughout all different stages of human neutrophilic granulopoiesis. This universal increase in neutrophil commitment is as expected given that both earlier proliferation and terminal differentiation of neutrophil progenitors were shown to be induced by G-CSF signaling^53^. The developmental stages of neutrophils in pre-Neu and Neu subsets were also confirmed with May-Grunwald Giemsa (MGG) staining (Fig. 3d).

**Figure 3.**
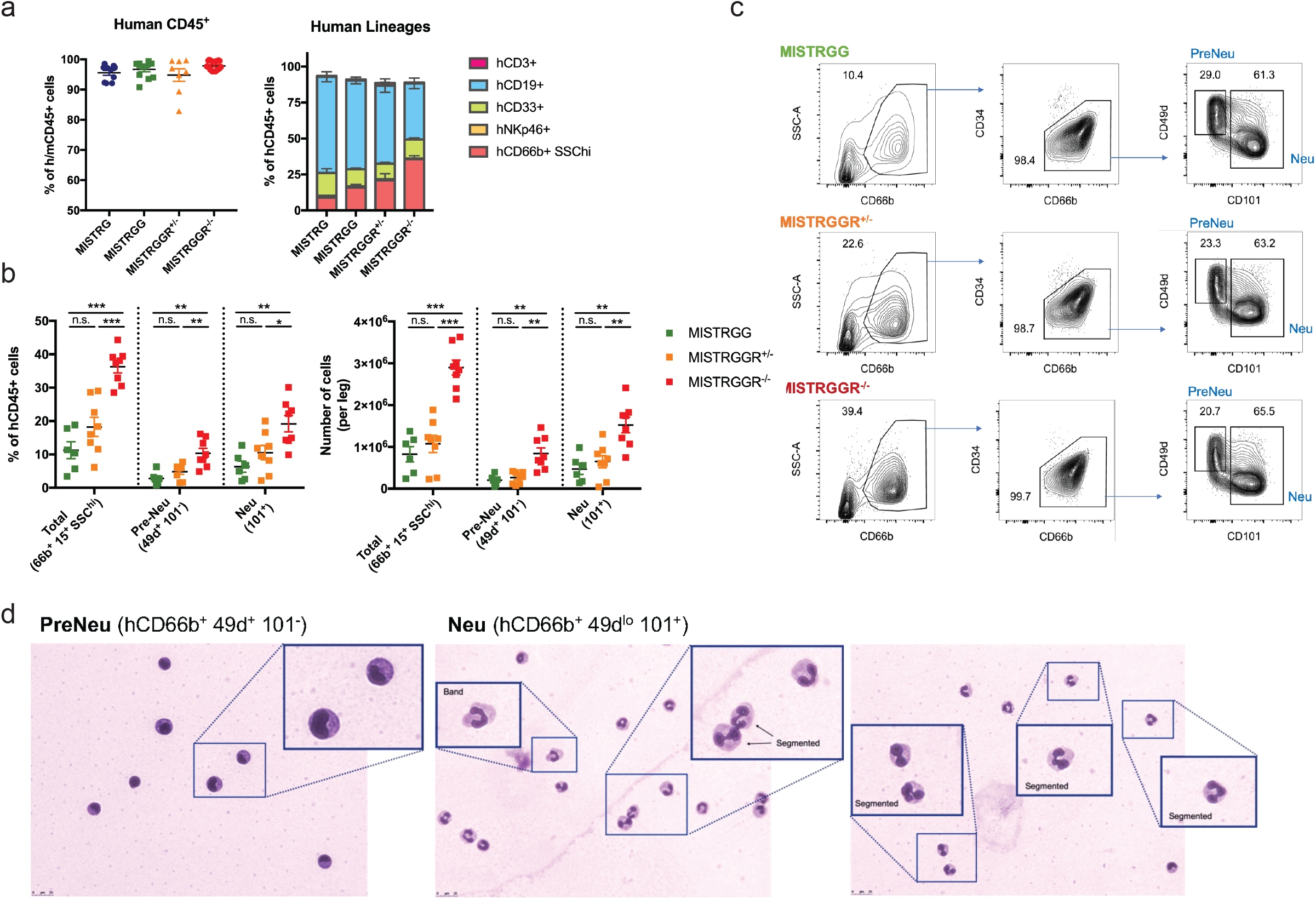
Enhanced bone marrow granulopoiesis in MISTRGGR-/- mice. Bone marrow cells were analyzed at 8 weeks post-en-graftment. (a) Frequencies of human hematopoietic cells (hCD45+) (left), human lineages (right) (MISTRG, n=5; MISTRGG, n=12; MISTRGGR+/-, n=8; MISTRGGR-/-, n=8). (b) Frequencies (left) and numbers (right) of human neutrophils (hCD66b+ SSChi), Pre-Neu (hCD49d+ CD101-) and Neu (hCD101+) in the bone marrow of MISTRGG, MISTRGGR+/-, and MISTRG-GR-/-. Data pooled from at least 2 independent experiments. (c) Representative flow cytometry gating of total human neutro-phils, Pre-Neu and Neu. (d) May-Grunwald Giemsa (MGG) staining of sorted bone marrow Pre-Neu (left) and Neu (right) cells. Enlarged boxes highlight various stages of human neutrophil development.

Next, we assessed the function of human neutrophils in MISTRGGR^-/-^ mice in peripheral tissues during inflammation. LPS nebulization provides an ideal tool to study recruitment and function of neutrophils in the lung. In this model, mice are exposed to aerosolized LPS, which initiates recruitment of neutrophils into the pulmonary vasculature, transendothelial migration into the lung interstitium, and subsequently transepithelial migration into the alveolar space^54^. Neutrophil accumulation in lung vasculature peaks at around 4 hours after nebulization, whereas the maximum migration into interstitial and alveolar spaces occurs at around 12-24 hours post-nebulization^54^. Engrafted 7 week-old MISTRG and MISTRGGR^-/-^ mice were exposed to aerosolized LPS and sacrificed at 24hr after nebulization. Bronchoalveolar lavage fluid (BALF), and lung tissues were harvested for flow cytometric assessment of infiltrating neutrophils and other immune cells (Fig. 4a). In order to differentiate immune cells residing in the lung vasculature from cells in the interstitium, we adopted a strategy of *in-vivo* labeling of circulating human immune cells^55^. Intravenous injection of anti-human CD45 allowed distinguishing between interstitial and intravascular neutrophils (Fig. 4b). At steady state, there were few human neutrophils in the lung of both MISTRGGR^-/-^ and MISTRG mice, although the numbers for MISTRGGR^-/-^ mice are significantly higher than MISTRG (Fig. 4c). However, upon LPS nebulization, there was a marked increase in human neutrophils in both intravenous and interstitial compartments in MISTRGGR^-/-^ mice but not in control MISTRG mice (Fig. 4c).

**Figure 4.**
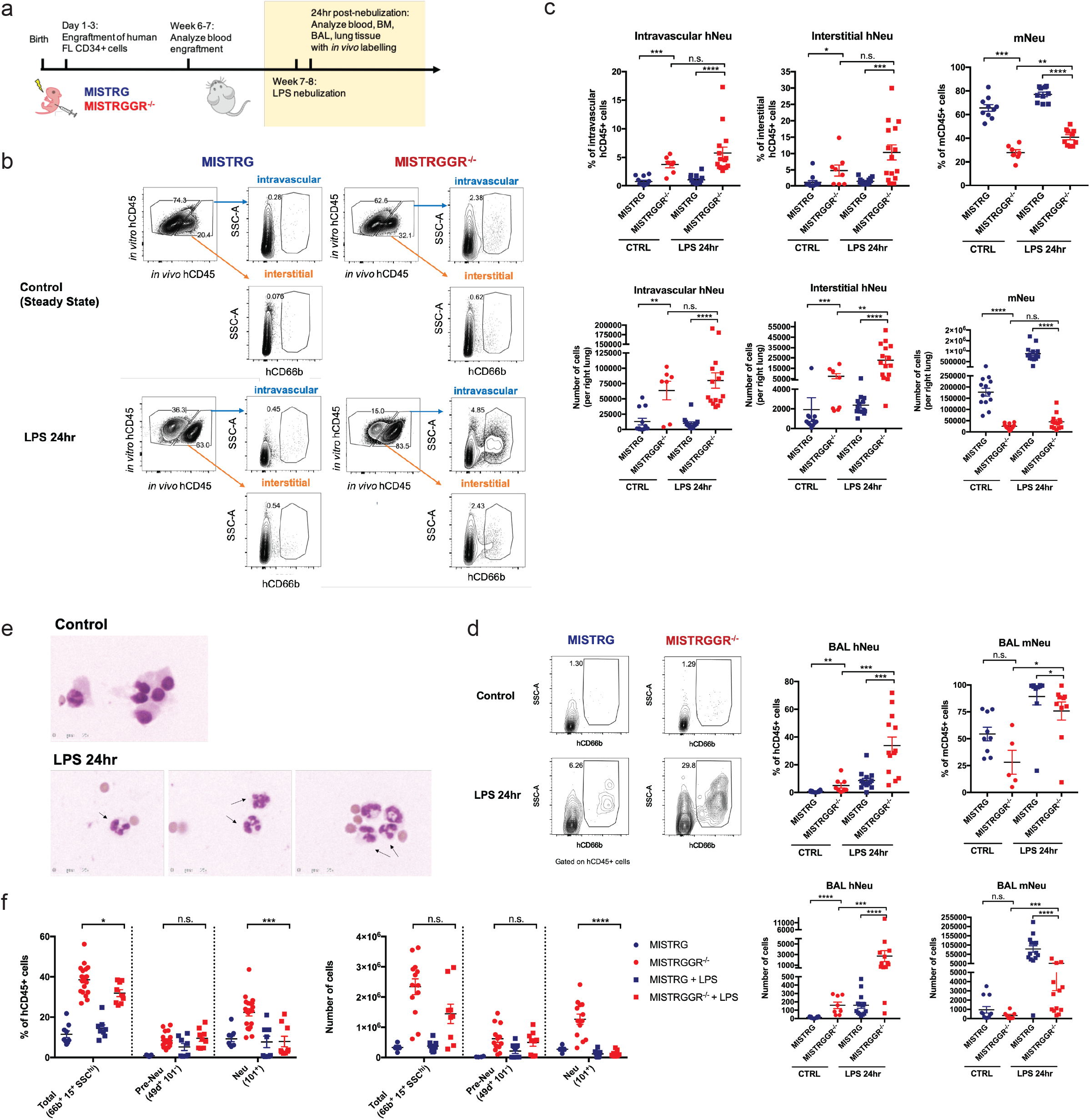
Robust human neutrophil recruitment to the lung upon inflammatory stimuli in MISTRGGR-/- mice. (a) Experimental timeline of LPS nebulization. 7-8 week-old engrafted mice were treated with LPS nebulization (12.5 mg administered over 15 min). Bronchoalveolar lavage fluid (BALF), and lung tissues were harvested at 24hr after nebulization. Prior to euthanasia, mice were injected with anti-human CD45 antibody intravenously to label circulating human cells. Interstitial neutrophils and intravascular neutrophils were distinguished by gating on in vivo versus in vitro hCD45 fluorochrome signals. (b, c) Representative flow cytometry plots (b) and quantifications (c) of intravascular and interstitial human neutrophils (hCD66b+ SSChi) and mouse neutrophils (Ly6G+) in lungs of MISTRG and MISTRGGR-/- mice at control steady state or 24hr after LPS nebulization. (d) Representative flow cytometry plots, frequencies and numbers of human and mouse neutrophils in BAL at control steady state or 24hr after LPS nebulization. Data pooled from at least 3 independent experiments. (e) May-Grunwald Giemsa (MGG) staining of cells isolated from BAL at 24hr after LPS nebulization. Arrows: human neutrophils. (f) Frequencies (top) and numbers (bottom) of human neutrophils (hCD66b+ SSChi), preNeu (hCD49d+ CD101-) and Neu (hCD101+) in the bone marrow of engrafted mice at steady state or 24hr after LPS nebulization. Data pooled from at least 2 independent experiments.

Although there was robust recruitment of mouse neutrophils in control MISTRG mice, human neutrophils failed to respond or expand (Fig. 4c). The recruitment of human neutrophils in MISTRGGR^-/-^ was even more dramatic in the BAL (Fig. 4d). The morphology of cells isolated from BAL were further confirmed by MGG staining. In contrast to steady state, where neutrophils are absent, BAL isolates from LPS-treated MISTRGGR^-/-^ mice are enriched with mature hyper-segmented human neutrophils (Fig. 4e). The accumulation of human neutrophils in the lung and BAL were accompanied by a significant reduction of CD101^+^ neutrophil (Neu) population in the bone marrow, suggesting a release of post-mitotic neutrophils in response to LPS stimuli (Fig. 4f). The levels of circulating human and mouse neutrophils and other human immune lineages in the blood remained unaffected by LPS nebulization (Fig. S2a, S4a, b, c). Human neutrophils in MISTRGGR^-/-^ mice are recruited into the pulmonary vasculature (Fig. 4b), migrate first into the lung interstitium (Fig. 4b, c), and then subsequently into the alveolar space (Fig. 4d). Overall, these observations demonstrate that human neutrophils generated in MISTRGGR^-/-^ mice are able to develop and respond robustly to inflammatory stimuli by homing to the sites of inflammation.

Finally, we tested the ability of human neutrophils to contain an infection, one of the major hallmarks of neutrophil function. *Pseudomonas aeruginosa*, a gram-negative bacterium, is one an opportunist pathogen that predominately infect neutropenic or immunocompromised individuals, such as cystic fibrosis patients, making it one of the leading causes of hospital-acquired pneumonia^25,56^. The clearance of *P. aeruginosa* during pulmonary infection is largely dependent on neutrophils and their intact machinery of intracellular killing following phagocytosis and extracellular killing following NET formation^25,57,58^. Given the relevance of *Pesudomonas* infections in human, we infected MISTRGGR^-/-^ and control MISTRG mice with *P. aeruginosa* intranasally (with a low and high dose of inoculation). In this experiment MISTRGGR^-/-^ and MISTRG mice were engrafted with fetal liver cells and were co-housed prior to infection to eliminate potential impacts of the microbiota. In order to assess the function of human neutrophils exclusive of mouse neutrophils, prior to infection we depleted the remaining mouse neutrophil compartment by administrating anti-mouse Ly6G antibody intravenously (Fig. 5a). The degree of depletion in blood and lung were validated by flow cytometry (Fig. S5a). Depletion of murine neutrophils by anti-Ly6G antibody also made MISTRG mice more susceptible to *P. aeruginosa* infection even at low doses of inoculation (Fig. S5b, S5c). Cell composition in lung vasculature, interstitium and BAL were assessed similarly by flow cytometry as previously stated (Fig. 4b). In all of these compartments, human neutrophils in MISTRGGR^-/-^ greatly outnumbered human neutrophils in MISTRG mice (Fig. 5b, 5c). MISTRGGR^-/-^ mice also had overall more human immune cells, but the composition of other immune cells except for granulocytes were comparable (Fig. 5d). Bacteria from lung and liver homogenates were plated and counted as an assessment of severity of infection and bacterial dissemination. MISTRGGR^-/-^ were able to better control *P. aeruginosa* infection as suggested by lower bacterial load in both lung and liver of MISTRGGR^-/-^ mice (Fig 5e). Therefore, these data show that human neutrophils generated in MISTRGGR^-/-^ mice are functional and capable of responding to and reducing bacterial burden. Overall, MISTRGGR^-/-^ mice enable development of functional, mature human neutrophils that are recruited during pulmonary infection to fulfill their essential role of fighting bacteria in-vivo.

**Figure 5.**
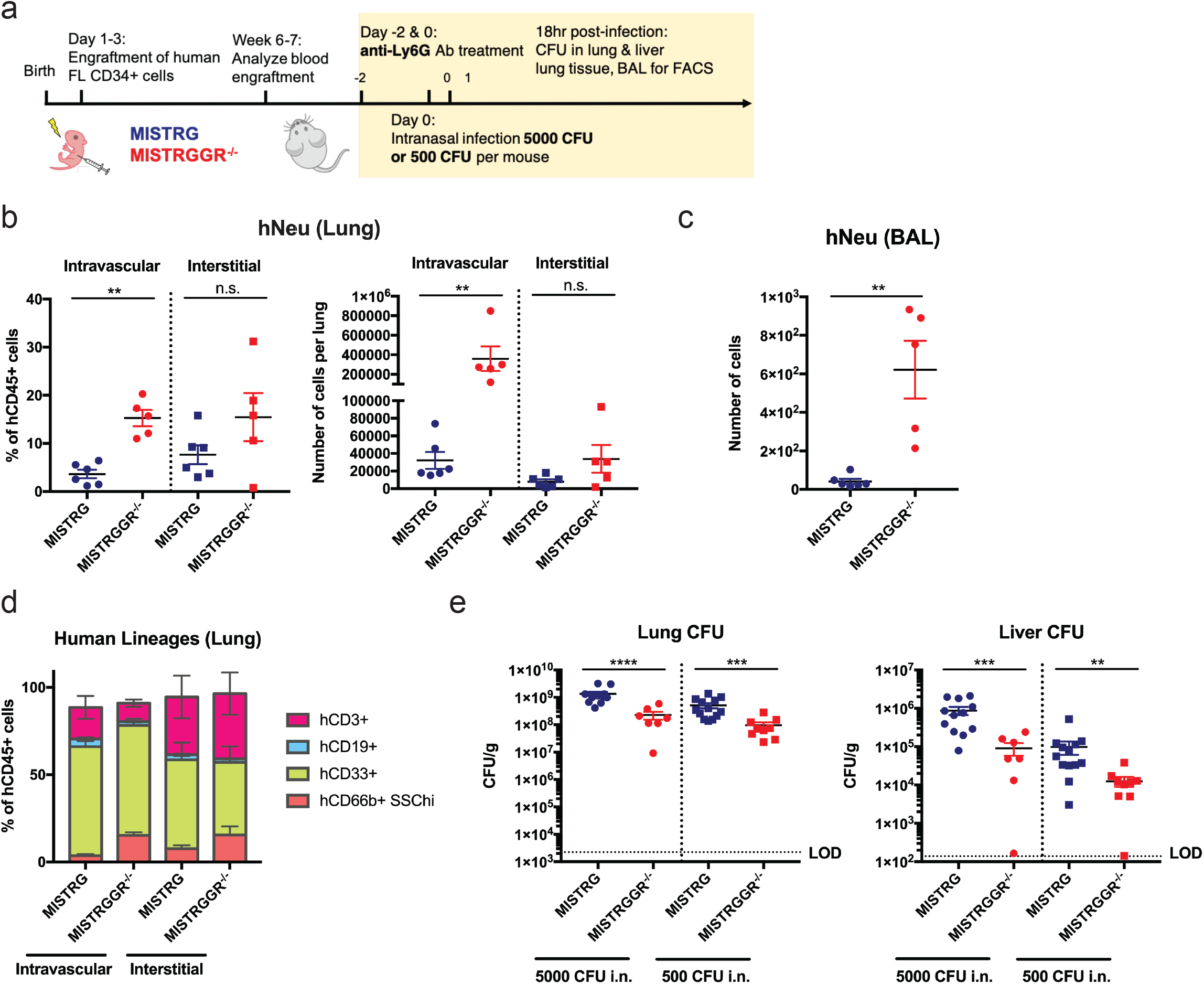
Human neutrophils in MISTRGGR-/- mice provide protection against Pseudomonas aeruginosa infection. (a) Experimental timeline of intranasal P. aeruginosa infection. Engrafted 7-8 wk-old MISTRG and MISTRGGR-/- mice were pre-treated with anti-mouse Ly6G antibody (1A8) (2 doses of 100 μg, i.v.) for depletion of murine neutrophils, and then intranasally infected with 500 or 5000 CFU of P. aeruginosa. Lungs and livers were harvested for CFU quantification and flow cytometric characterization at 18hr after infection. (b, c) Frequen-cies and numbers of human neutrophils in the lung (b) and BAL (c) of infected mice (500 CFU, 18hr p.i.). (d) Frequencies of human lineages (MISTRG, n=5; MISTRGGR-/-, n=6) in intravascular and interstitial lung post-in-fection (500 CFU, 18hr p.i.). (e) Bacterial load in lung (left) and liver (right) at 18hr after infection. Data pooled from at least 3 independent experiments.

## Discussion

Humanized mouse models have greatly expanded our understanding of how the human immune system works and notably, have aided in translating findings in mice to humans. However, until now, there were no humanized mouse models that supported mature functional neutrophils which are a cornerstone of the innate immune response. MISTRGGR^-/-^ mice described here overcome this major limitation in current generation humanized models by greatly improve human neutrophil numbers in blood and peripheral tissues. Importantly, this model allows for the first-time functional integration of human neutrophil function in vivo and greatly expands the utility of humanized mice for studying complex human disease.

In our system, humanization of G-CSF alone did not improve human neutrophils. Results achieved by deletion of G-CSFR, receptor that binds to G-CSF argues that this is due to cross-reactivity of mouse G-CSFR with human G-CSF, which display a relatively high homology to mouse G-CSF^46^. Given that murine granulocytes and precursors are still in vast majority during the early weeks of engraftment, the presence of murine cells will likely have a competitive advantage over the human G-CSF and occupy niche space for granulopoiesis. Therefore, G-CSFR deletion was necessary to limit murine granulopoiesis and to allow human granulocyte development. This model still exhibits deficiencies. Although MISTRGGR^-/-^ mice have significantly fewer mouse neutrophils, G-CSFR deficiency did not completely eliminate mouse neutrophils, opening up possibilities for further improvement of the existing system. Additionally, at steady state, human neutrophil numbers seen in blood of MISTRGGR^-/-^ mice are still not comparable to the high proportions and numbers of neutrophils seen in human blood. We have confirmed a pool of post-mitotic mature human neutrophils in the bone marrow of engrafted MISTRGGR^-/-^ mice (Fig 3d), and yet these neutrophils display lower CD10 expression, a marker for mature human bone marrow neutrophils^52^, as compared to mature neutrophils isolated from adult human bone marrow (Fig S3b, c, Ref. ^52^). Nevertheless, our model is a major step forward to understanding why and will allow more mechanistic studies to be performed to understand how this could be achieved and what additional unknown factors are required for human neutrophil development.

Although human neutrophils effectively home to lungs and are functional in reducing bacterial burden during *P. aeruginosa* infection, the presence of human neutrophils does not completely clear the infection. MISTRG mice, without depletion of murine neutrophils have a lower bacterial load compared with engrafted MISTRGGR^-/-^ mice. It is unclear whether it is just a quantitative deficiency as there are still fewer human neutrophils in engrafted MISTRGGR^-/-^ than there are mouse neutrophils in MISTRG mice (Fig 5b, S5c). Other qualitative functions of infiltrating human neutrophils (e.g., NETs formation) during infection also remain elusive, some of which could potentially require further support in the humanized host.

A human immune system that supports mature neutrophils can be combined with various disease models to study human specific contribution of neutrophils during disease. Epidemics of drug-resistant and opportunist pathogens have emphasized the need to further investigate the interactions between human neutrophils and pathogens. *Candida auris*, a recently-emerged fungal pathogen posing global public health threats, has been shown to be capable of resisting human neutrophil killing by avoiding effective engagement, phagocytosis, as well as inhibiting the formation of NETs^26^. Moreover, in contrast to the inability of human neutrophils to recognize and kill the pathogen, murine neutrophils appear to be able to control *C. auris* infection^59-61^, suggesting a species-specific mechanism by which *C. auris* are able to resist human neutrophil killing. Given that our knowledge of neutrophil responses to infection has been largely derived from animal models and *in vitro* studies, understanding the *in vivo* action of human neutrophils in various bacterial and fungal infections remains critical. Here, we have demonstrated the potential to use MISTRGGR^-/-^ mice to study the interactions between human neutrophils and respiratory pathogens in the native lung environment.

In addition to antimicrobial properties of neutrophils, there has been a growing appreciation of neutrophils in tumorigenesis. However, the current understanding of their roles still remains in debate, as tumor-associated neutrophils, upon activation display both pro and anti-tumor properties^23,32,33,62-67^. It remains unclear whether differential programs for neutrophils (similar to M1 and M2 for macrophages) in tumor environment truly exists, given that the heterogeneity observed could be due to differential states of activation and maturation^68,69^. To date, the majority of experimental approaches for studying neutrophils in tumor tissues have been limited to mouse models as well as correlative studies of human neutrophils in various patient tumors. Thus, our model provides a unique possibility to co-transplant the human immune system and human tumors, which will allow the study of the role of neutrophils in tumor evolution and help to identify potential therapy targets.

Our humanized model that targets the cytokine-receptor pair of G-CSF significantly improves functional human neutrophil presence in periphery but also highlights the importance of multiple strategies that are still needed to fully recapitulate functional human immune response. MISTRGGR^-/-^ mice are an important and necessary step forward in pushing the next generation of humanized mouse systems and provide a unique tool to study human neutrophils *in vivo* in presence of a more complete human immune system. Together, this in turn should enable the application of humanized mice to unexplored and wider areas of translational research.

## Acknowledgements

We thank G. Yancopoulos, D. Valenzuela, A. Murphy, and W. Auerbach at Regeneron Pharmaceuticals who generated, in collaboration with our groups, the individual knock-in alleles combined in MISTRG; Jon Alderman, Caroline Lieber, and Elizabeth Hughes-Picard for administrative assistance; Carla Weibel, Patricia Ranney, Cynthia Hughes for mouse colony management; Judith Stein and Linda Evangelisti for mouse engineering; David Urbanos for human CD34+ cell isolation; Yuanbin Song and Xiaoying Fu for the help with cytospin and discussion; Caterina Di Pietro for assistance with LPS nebulization; C. Lieber for manuscript submission and Michael Chiorazzi, Ian Odell, Dietmar Herndler-Brandstetter and Elizabeth Eynon for discussion. E.S. is a HHMI Fellow of the Damon Runyon Cancer Research Foundation (DRG-2316-18). This work was funded by the Howard Hughes Medical Institute, NIH 1U54DK106857 and Yale Cooperative Hematology Specialized Core Center (R.A.F.).

## Authorship Contributions

Y.Z. and E.S. designed and performed experiments, analyzed the data, and wrote the manuscript; A.G.S., and H.H.O. performed experiments and edited the manuscript; J.A. generated the human G-CSF knock-in mice and edited the manuscript; R.J., E.M.B., S.H. and L.S. provided valuable input for this study; R.A.F. conceived the project, supervised its participants, interpreted its results, and edited the manuscript.

## References

1 Mestas, J. & Hughes, C. C. Of mice and not men: differences between mouse and human immunology. J Immunol 172, 2731–2738, doi:10.4049/jimmunol.172.5.2731 (2004).

2 Gibbons, D. L. & Spencer, J. Mouse and human intestinal immunity: same ballpark, different players; different rules, same score. Mucosal Immunol 4, 148–157, doi:10.1038/mi.2010.85 (2011).

3 Rongvaux, A. et al. Human hemato-lymphoid system mice: current use and future potential for medicine. Annu Rev Immunol 31, 635–674, doi:10.1146/annurev-immunol-032712-095921 (2013).

4 Zschaler, J., Schlorke, D. & Arnhold, J. Differences in innate immune response between man and mouse. Crit Rev Immunol 34, 433–454 (2014).

5 Shay, T. et al. Conservation and divergence in the transcriptional programs of the human and mouse immune systems. Proc Natl Acad Sci U S A 110, 2946–2951, doi:10.1073/pnas.1222738110 (2013).

6 Ng, L. G., Ostuni, R. & Hidalgo, A. Heterogeneity of neutrophils. Nat Rev Immunol 19, 255–265, doi:10.1038/s41577-019-0141-8 (2019).

7 Lok, L. S. C. et al. Phenotypically distinct neutrophils patrol uninfected human and mouse lymph nodes. Proc Natl Acad Sci U S A 116, 19083–19089, doi:10.1073/pnas.1905054116 (2019).

8 Casanova-Acebes, M. et al. Neutrophils instruct homeostatic and pathological states in naive tissues. J Exp Med 215, 2778–2795, doi:10.1084/jem.20181468 (2018).

9 Kolaczkowska, E. & Kubes, P. Neutrophil recruitment and function in health and inflammation. Nat Rev Immunol 13, 159–175, doi:10.1038/nri3399 (2013).

10 Nauseef, W. M. & Borregaard, N. Neutrophils at work. Nat Immunol 15, 602–611, doi:10.1038/ni.2921 (2014).

11 Scapini, P. et al. The neutrophil as a cellular source of chemokines. Immunol Rev 177, 195–203, doi:10.1034/j.1600-065x.2000.17706.x (2000).

12 Zollner, O. et al. L-selectin from human, but not from mouse neutrophils binds directly to E-selectin. J Cell Biol 136, 707–716, doi:10.1083/jcb.136.3.707 (1997).

13 Aleyd, E., Heineke, M. H. & van Egmond, M. The era of the immunoglobulin A Fc receptor FcalphaRI; its function and potential as target in disease. Immunol Rev 268, 123–138, doi:10.1111/imr.12337 (2015).

14 Hajjar, E., Broemstrup, T., Kantari, C., Witko-Sarsat, V. & Reuter, N. Structures of human proteinase 3 and neutrophil elastase--so similar yet so different. FEBS J 277, 2238–2254, doi:10.1111/j.1742-4658.2010.07659.x (2010).

15 Otten, M. A. et al. Immature neutrophils mediate tumor cell killing via IgA but not IgG Fc receptors. J Immunol 174, 5472–5480, doi:10.4049/jimmunol.174.9.5472 (2005).

16 Tecchio, C., Micheletti, A. & Cassatella, M. A. Neutrophil-derived cytokines: facts beyond expression. Front Immunol 5, 508, doi:10.3389/fimmu.2014.00508 (2014).

17 Tamassia, N. et al. Cutting edge: An inactive chromatin configuration at the IL-10 locus in human neutrophils. J Immunol 190, 1921–1925, doi:10.4049/jimmunol.1203022 (2013).

18 Condliffe, A. M. et al. Sequential activation of class IB and class IA PI3K is important for the primed respiratory burst of human but not murine neutrophils. Blood 106, 1432–1440, doi:10.1182/blood-2005-03-0944 (2005).

19 Bagaitkar, J., Matute, J. D., Austin, A., Arias, A. A. & Dinauer, M. C. Activation of neutrophil respiratory burst by fungal particles requires phosphatidylinositol 3-phosphate binding to p40(phox) in humans but not in mice. Blood 120, 3385–3387, doi:10.1182/blood-2012-07-445619 (2012).

20 Ganz, T. Defensins: antimicrobial peptides of innate immunity. Nat Rev Immunol 3, 710–720, doi:10.1038/nri1180 (2003).

21 Eisenhauer, P. B. & Lehrer, R. I. Mouse neutrophils lack defensins. Infect Immun 60, 3446–3447 (1992).

22 Rausch, P. G. & Moore, T. G. Granule enzymes of polymorphonuclear neutrophils: A phylogenetic comparison. Blood 46, 913–919 (1975).

23 Eruslanov, E. B., Singhal, S. & Albelda, S. M. Mouse versus Human Neutrophils in Cancer: A Major Knowledge Gap. Trends Cancer 3, 149–160, doi:10.1016/j.trecan.2016.12.006 (2017).

24 Witter, A. R., Okunnu, B. M. & Berg, R. E. The Essential Role of Neutrophils during Infection with the Intracellular Bacterial Pathogen Listeria monocytogenes. J Immunol 197, 1557–1565, doi:10.4049/jimmunol.1600599 (2016).

25 Rada, B. Interactions between Neutrophils and Pseudomonas aeruginosa in Cystic Fibrosis. Pathogens 6, doi:10.3390/pathogens6010010 (2017).

26 Johnson, C. J., Davis, J. M., Huttenlocher, A., Kernien, J. F. & Nett, J. E. Emerging Fungal Pathogen Candida auris Evades Neutrophil Attack. MBio 9, doi:10.1128/mBio.01403-18 (2018).

27 Jones-Nelson, O. et al. The Neutrophilic Response to Pseudomonas Damages the Airway Barrier, Promoting Infection by Klebsiella pneumoniae. Am J Respir Cell Mol Biol 59, 745–756, doi:10.1165/rcmb.2018-0107OC (2018).

28 Lieschke, G. J. et al. Mice lacking granulocyte colony-stimulating factor have chronic neutropenia, granulocyte and macrophage progenitor cell deficiency, and impaired neutrophil mobilization. Blood 84, 1737–1746 (1994).

29 Triot, A. et al. Inherited biallelic CSF3R mutations in severe congenital neutropenia. Blood 123, 3811–3817, doi:10.1182/blood-2013-11-535419 (2014).

30 Morstyn, G. et al. Effect of granulocyte colony stimulating factor on neutropenia induced by cytotoxic chemotherapy. Lancet 1, 667–672, doi:10.1016/s0140-6736(88)91475-4 (1988).

31 Russell, A. R., Davies, E. G., Ball, S. E. & Gordon-Smith, E. Granulocyte colony stimulating factor treatment for neonatal neutropenia. Arch Dis Child Fetal Neonatal Ed 72, F53–54, doi:10.1136/fn.72.1.f53 (1995).

32 Fridlender, Z. G. et al. Polarization of tumor-associated neutrophil phenotype by TGF-beta: “N1” versus “N2” TAN. Cancer Cell 16, 183–194, doi:10.1016/j.ccr.2009.06.017 (2009).

33 Nozawa, H., Chiu, C. & Hanahan, D. Infiltrating neutrophils mediate the initial angiogenic switch in a mouse model of multistage carcinogenesis. Proc Natl Acad Sci U S A 103, 12493–12498, doi:10.1073/pnas.0601807103 (2006).

34 Puga, I. et al. B cell-helper neutrophils stimulate the diversification and production of immunoglobulin in the marginal zone of the spleen. Nat Immunol 13, 170–180, doi:10.1038/ni.2194 (2011).

35 Rongvaux, A. et al. Development and function of human innate immune cells in a humanized mouse model. Nat Biotechnol 32, 364–372, doi:10.1038/nbt.2858 (2014).

36 Fine, N., Barzilay, O. & Glogauer, M. Analysis of Human and Mouse Neutrophil Phagocytosis by Flow Cytometry. Methods Mol Biol 1519, 17–24, doi:10.1007/978-1-4939-6581-6_2 (2017).

37 Taylor, A. et al. SRF is required for neutrophil migration in response to inflammation. Blood 123, 3027–3036, doi:10.1182/blood-2013-06-507582 (2014).

38 Sun, F., Xiao, G. & Qu, Z. Murine Bronchoalveolar Lavage. Bio Protoc 7, doi:10.21769/BioProtoc.2287 (2017).

39 Christopher, M. J. & Link, D. C. Regulation of neutrophil homeostasis. Curr Opin Hematol 14, 3–8 (2007).

40 Panopoulos, A. D. & Watowich, S. S. Granulocyte colony-stimulating factor: molecular mechanisms of action during steady state and ‘emergency’ hematopoiesis. Cytokine 42, 277–288, doi:10.1016/j.cyto.2008.03.002 (2008).

41 Semerad, C. L., Liu, F., Gregory, A. D., Stumpf, K. & Link, D. C. G-CSF is an essential regulator of neutrophil trafficking from the bone marrow to the blood. Immunity 17, 413–423, doi:10.1016/s1074-7613(02)00424-7 (2002).

42 Strydom, N. & Rankin, S. M. Regulation of circulating neutrophil numbers under homeostasis and in disease. J Innate Immun 5, 304–314, doi:10.1159/000350282 (2013).

43 Boettcher, S. et al. Endothelial cells translate pathogen signals into G-CSF-driven emergency granulopoiesis. Blood 124, 1393–1403, doi:10.1182/blood-2014-04-570762 (2014).

44 Manz, M. G. & Boettcher, S. Emergency granulopoiesis. Nat Rev Immunol 14, 302–314, doi:10.1038/nri3660 (2014).

45 Nicola, N. A. Granulocyte colony-stimulating factor and differentiation-induction in myeloid leukemic cells. Int J Cell Cloning 5, 1–15, doi:10.1002/stem.5530050102 (1987).

46 Nicola, N. A., Begley, C. G. & Metcalf, D. Identification of the human analogue of a regulator that induces differentiation in murine leukaemic cells. Nature 314, 625–628, doi:10.1038/314625a0 (1985).

47 Ran, F. A. et al. Genome engineering using the CRISPR-Cas9 system. Nat Protoc 8, 2281–2308, doi:10.1038/nprot.2013.143 (2013).

48 Bajrami, B. et al. G-CSF maintains controlled neutrophil mobilization during acute inflammation by negatively regulating CXCR2 signaling. J Exp Med 213, 1999–2018, doi:10.1084/jem.20160393 (2016).

49 Zsebo, K. M. et al. Recombinant human granulocyte colony stimulating factor: molecular and biological characterization. Immunobiology 172, 175–184, doi:10.1016/S0171-2985(86)80097-3 (1986).

50 Liu, F., Wu, H. Y., Wesselschmidt, R., Kornaga, T. & Link, D. C. Impaired production and increased apoptosis of neutrophils in granulocyte colony-stimulating factor receptor-deficient mice. Immunity 5, 491–501, doi:10.1016/s1074-7613(00)80504-x (1996).

51 Pillay, J., Tak, T., Kamp, V. M. & Koenderman, L. Immune suppression by neutrophils and granulocytic myeloid-derived suppressor cells: similarities and differences. Cell Mol Life Sci 70, 3813–3827, doi:10.1007/s00018-013-1286-4 (2013).

52 Evrard, M. et al. Developmental Analysis of Bone Marrow Neutrophils Reveals Populations Specialized in Expansion, Trafficking, and Effector Functions. Immunity 48, 364–379 e368, doi:10.1016/j.immuni.2018.02.002 (2018).

53 Liongue, C., Wright, C., Russell, A. P. & Ward, A. C. Granulocyte colony-stimulating factor receptor: stimulating granulopoiesis and much more. Int J Biochem Cell Biol 41, 2372–2375, doi:10.1016/j.biocel.2009.08.011 (2009).

54 Reutershan, J., Basit, A., Galkina, E. V. & Ley, K. Sequential recruitment of neutrophils into lung and bronchoalveolar lavage fluid in LPS-induced acute lung injury. Am J Physiol Lung Cell Mol Physiol 289, L807–815, doi:10.1152/ajplung.00477.2004 (2005).

55 Solis, A. G. et al. Mechanosensation of cyclical force by PIEZO1 is essential for innate immunity. Nature 573, 69–74, doi:10.1038/s41586-019-1485-8 (2019).

56 Mandal, P. K. et al. Micro-organisms Associated with Febrile Neutropenia in Patients with Haematological Malignancies in a Tertiary Care Hospital in Eastern India. Indian J Hematol Blood Transfus 31, 46–50, doi:10.1007/s12288-014-0393-1 (2015).

57 Lavoie, E. G., Wangdi, T. & Kazmierczak, B. I. Innate immune responses to Pseudomonas aeruginosa infection. Microbes Infect 13, 1133–1145, doi:10.1016/j.micinf.2011.07.011 (2011).

58 Young, R. L. et al. Neutrophil extracellular trap (NET)-mediated killing of Pseudomonas aeruginosa: evidence of acquired resistance within the CF airway, independent of CFTR. PLoS One 6, e23637, doi:10.1371/journal.pone.0023637 (2011).

59 Ferwerda, B. et al. Human dectin-1 deficiency and mucocutaneous fungal infections. N Engl J Med 361, 1760–1767, doi:10.1056/NEJMoa0901053 (2009).

60 Li, X. et al. The beta-glucan receptor Dectin-1 activates the integrin Mac-1 in neutrophils via Vav protein signaling to promote Candida albicans clearance. Cell Host Microbe 10, 603–615, doi:10.1016/j.chom.2011.10.009 (2011).

61 Ben-Ami, R. et al. Multidrug-Resistant Candida haemulonii and C. auris, Tel Aviv, Israel. Emerg Infect Dis 23, doi:10.3201/eid2302.161486 (2017).

62 Albanesi, M. et al. Neutrophils mediate antibody-induced antitumor effects in mice. Blood 122, 3160–3164, doi:10.1182/blood-2013-04-497446 (2013).

63 Houghton, A. M. et al. Neutrophil elastase-mediated degradation of IRS-1 accelerates lung tumor growth. Nat Med 16, 219–223, doi:10.1038/nm.2084 (2010).

64 Granot, Z. et al. Tumor entrained neutrophils inhibit seeding in the premetastatic lung. Cancer Cell 20, 300–314, doi:10.1016/j.ccr.2011.08.012 (2011).

65 MET Promotes Antitumor Neutrophil Recruitment and Cytotoxicity. Cancer Discov 5, 689, doi:10.1158/2159-8290.CD-RW2015-097 (2015).

66 Blaisdell, A. et al. Neutrophils Oppose Uterine Epithelial Carcinogenesis via Debridement of Hypoxic Tumor Cells. Cancer Cell 28, 785–799, doi:10.1016/j.ccell.2015.11.005 (2015).

67 Liu, Y. et al. CD11b+Ly6G+ cells inhibit tumor growth by suppressing IL-17 production at early stages of tumorigenesis. Oncoimmunology 5, e1061175, doi:10.1080/2162402X.2015.1061175 (2016).

68 Adrover, J. M., Nicolas-Avila, J. A. & Hidalgo, A. Aging: A Temporal Dimension for Neutrophils. Trends Immunol 37, 334–345, doi:10.1016/j.it.2016.03.005 (2016).

69 Nicolas-Avila, J. A., Adrover, J. M. & Hidalgo, A. Neutrophils in Homeostasis, Immunity, and Cancer. Immunity 46, 15–28, doi:10.1016/j.immuni.2016.12.012 (2017).

