## Supplemental Data for "Human neutrophil development and functionality are enabled in a humanized mouse model"

### Materials and Methods

#### Generation of MISTRGGR mice.

The generation of GCSF knock-in mice was performed on a MISTRG background with CRISPR guide designed to target region (chr11:98,701,464-98,701,486) in intron 1 of mouse *G-CSF*. Guide sequence: GGTCTGAGGCACTTGTTATCCGG. A construct containing human *GCSF* ORF (1485bp) flanked by two homologous arms were provided. The resulting founding mouse contains the entire human ORF with murine 5' and 3' UTRs. The knock-out of *G-CSFR* were conducted on MISTRGG mice with CRISPR guides designed to target both the ATG start codon and the beginning of 3' UTR. Guide #1 sequences: GCTGGCAAAATGGTAGGGCTGGG. Guide #2 sequences: GAAATTCTGGCTGGACGTGGTGG. The resulting founding mouse contains a deletion of 16798 bp (mChr4:126,027,350-126,044,148), starting immediately after start codon and 310 bp into 3' UTR of G-CSFR. All mice were maintained under specific pathogen-free conditions with continuous treatment with enrofloxacin in the drinking water (Baytril, 0.27 mg/ml). All animal experimentations were performed in compliance with Yale Institutional Animal Care and Use Committee protocols.

#### Gene Expression Analysis.

Total RNA was extracted from tissues using TRIzol reagent (Thermo Fisher) and the DirectZol RNA Miniprep Kit (Zymo Research). cDNA synthesis was performed using SuperScript III Reverse Transcriptase (Thermo Fisher). Quantitative reverse transcription PCR (qRT-PCR) was performed using a CFX96™ Real-Time PCR System (Bio-Rad) and iTaq Universal Probes Supermix (Bio-Rad). Sequence-specific oligonucleotide primers were purchased from Sigma-Aldrich. The following primers were used: human CSF3 (Hs00738432_g1), mouse Csf3 (Mm00438334_m1), mouse Csf3r (Mm00432735_m1). Expression values were calculated using the comparative threshold cycle method and normalized to mouse Hprt (Mm00446968_m1).

#### Characterization of human immune cells by flow cytometry.

All mice were analyzed at approximately 6-8 weeks post-engraftment. Blood was collected retro-orbitally, stained by an antibody cocktail at room temperature for 20 min and then treated with RBC lysis and fixation buffer (Biolegend) before analysis. Femurs and tibias were dissected and flushed with PBS supplemented with 2% FBS and 1mM EDTA through a 30-gauge needle (BD) to form single-cell suspensions, which were subsequently treated with ammonium-chloride-potassium (ACK) lysis buffer to eliminate RBCs. Lung tissues were chopped into small pieces, digested in RPMI1640 medium supplemented with collagenase D (1mg/ml) for 35 min at 37 °C, and filtered through 70um cell strainer to form single cell suspensions. Cell suspensions of BMs, lungs and BALs were stained with antibody cocktail on ice for 20 min, washed with PBS (2% FBS, 1mM EDTA) and fixed with Foxp3 / Transcription Factor Fixation Buffer (eBioscience) overnight.

Antibodies against the following antigens were used at final concentration of 0.5 µg/ml: Mouse antigens: CD45 (Clone: 30-F11), Ly6G (1A8), Ly6C (HK1.4); Human antigens: CD45 (HI30), CD3 (HIT3a), CD10 (HI10a), CD14 (HCD14), CD15 (W6D3), CD16 (3G8), CD19 (HIB19), CD33 (WM53), CD34 (561), CD49d (9F10), CD66b (G10F5), CD101 (BB27), NKp46 (9E2). All antibodies were obtained from Biolegend, unless otherwise specified. Data were acquired with FACSDiva on an LSRII flow cytometer (BD Biosciences) and analyzed with FlowJo software.

**Cell morphology.**
Cells were cytospun onto slides at 200 rpm for 5 min (~100,000 cells/slide) and stained with May-Grunwald Giemsa (MGG, Sigma). Blood smears were performed using standard methods and stained with MGG.

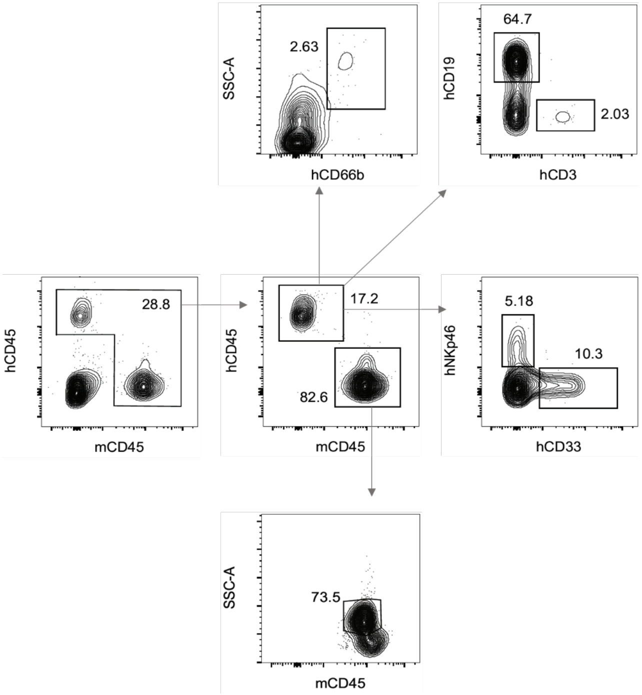


Figure S1. Characterization of human immune lineages in engrafted MISTRG mice. Representative flow plots for the gating strategy for human immune cell lineages in the blood are shown. Live single cells were first gated for a combined human(h) and mouse(m) CD45+, gate and then hCD45+ and mCD45+ respectively. hCD45+ cells were subsequently gated for hCD3+ (T cells), hCD19+ (B cells), hCD33+ (myeloid cells/monocytes), hCD66b+ SSChi (neutrophils), and hNKp46+ (NK cells). mCD45+ population were gated for SSChi (neutrophils). In later experiments, mouse neutrophils were also identified by mLy6G+, or Ly6Cmid SSChi.

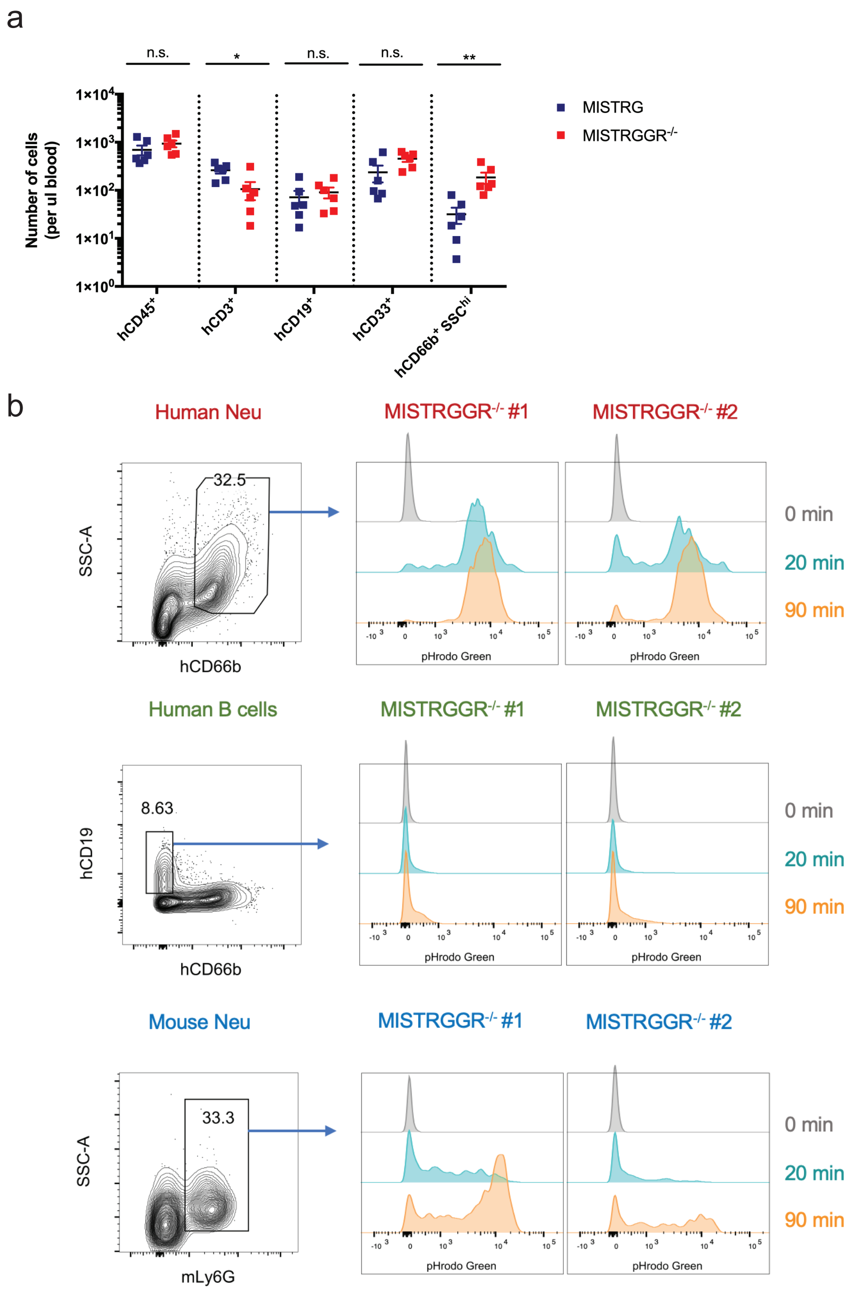


Figure S2. Characterization of human blood immune cells in engrafted MISTRG and MISTRGGR-/- mice. (a) Quantifications of human immune lineages in blood at week 7 post-engraftment. Data pooled from at least 2 independent experiments. All mice were irradiated with 150 Rads and intra-hepatically injected with 20,000 human fetal liver CD34+ cells at 1-3 days after birth. (b) Representative flow cytometry analysis of phagocytosis by reconstituted human neutrophils (top), human B cells (middle) and mouse neutrophils (bottom) from the blood of engrafted MISTRGGR-/- mice. Blood samples were incubated with pHrodo™ Green E. coli BioParticles® Conjugate (10ug per 50ul blood) for the indicated times (0, 20 and 90 min) and fluorescent signals were analyzed by flow cytometry. Data were represenative of at least 2 independent experiments.

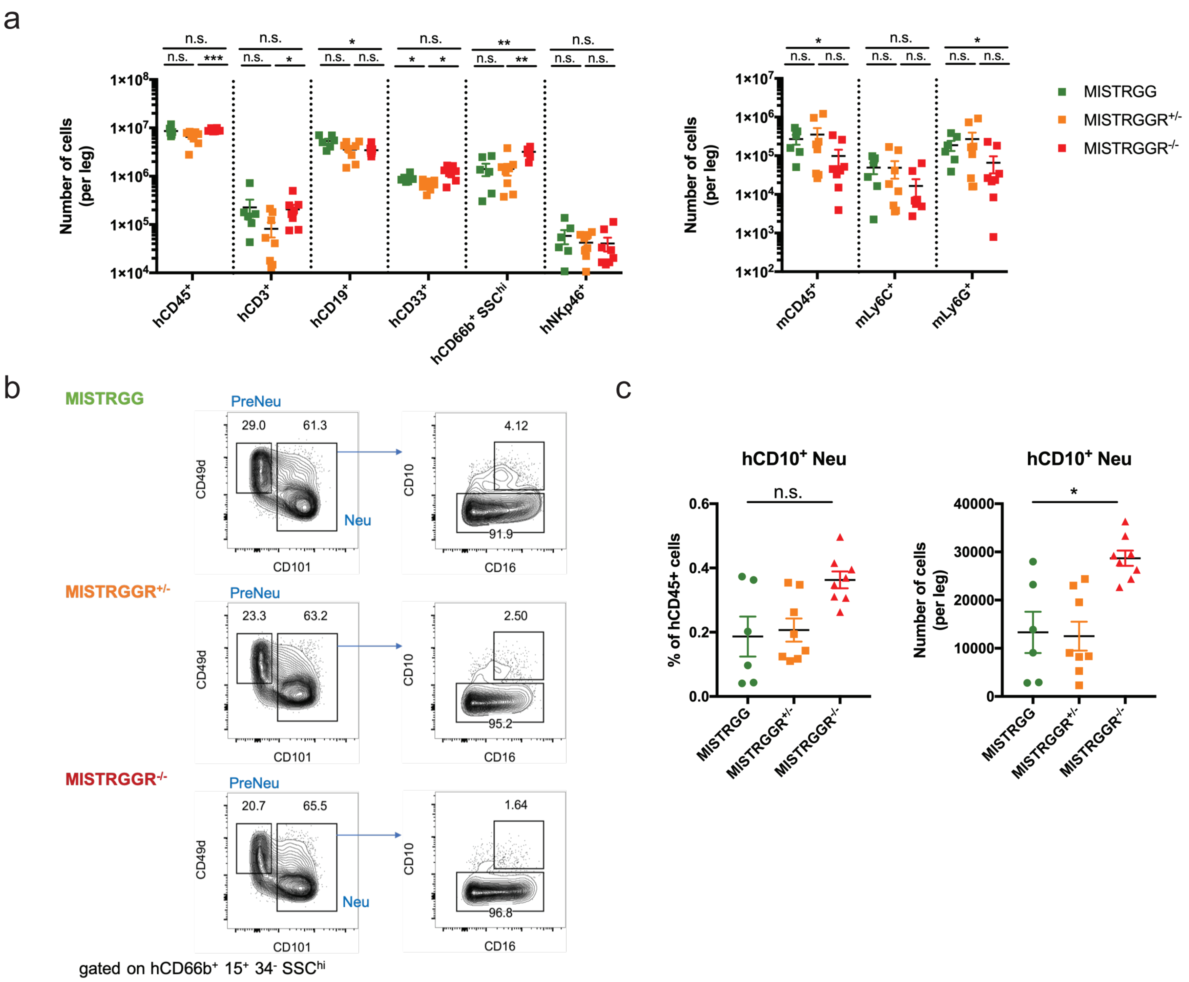


Figure S3. Characterization of bone marrow engraftment in MISTRGG, MISTRGGR+/- and MISTRGGR-/- mice. (a) Quantifications of human (left) and mouse (right) immune lineages in engrafted bone marrow at week 8 post-engraftment. (b) Representative flow cytometry plots of CD10 expression in Neu subset. (c) Frequencies (left) and numbers (right) of CD10+ Neu. Data pooled from at least 2 independent experiments. All mice were irradiated with 150 Rads and intra-hepatically injected with 20,000 human fetal liver CD34+ cells at 1-3 days after birth.

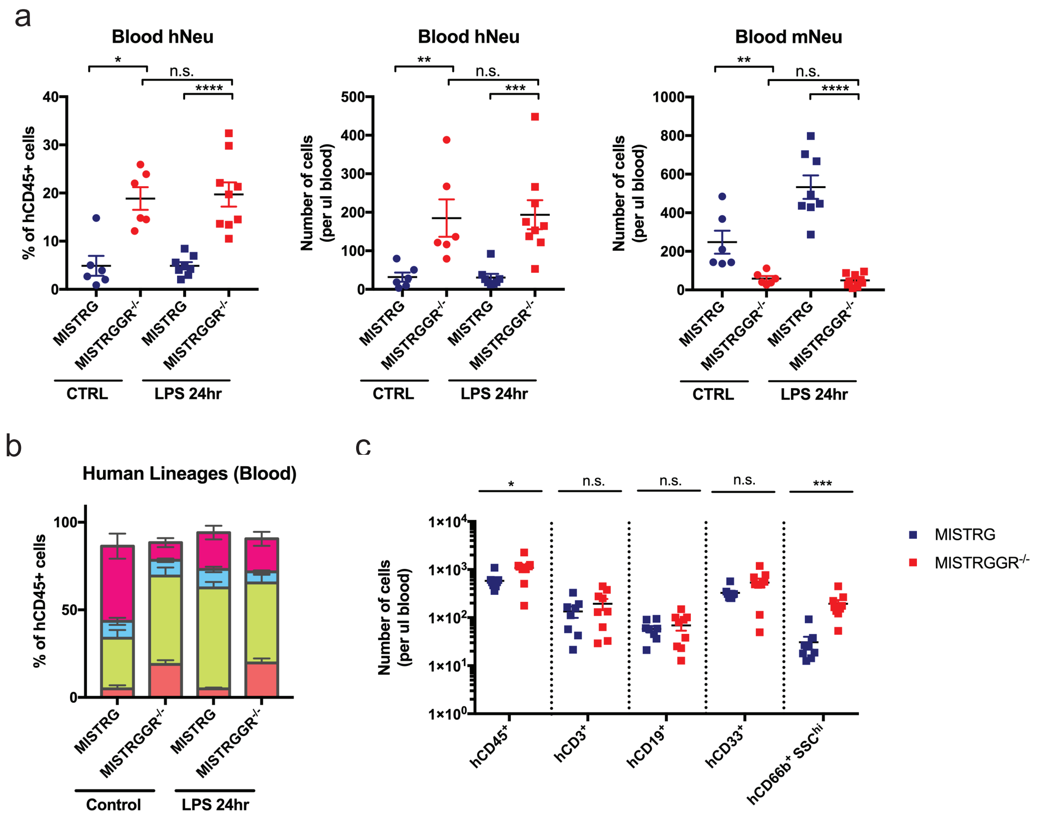


Figure S4. Characterization of human immune cells in the blood of engrafted MISTRG and MISTRGGR-/- mice in response to LPS nebulization. (a) Quantifications of human and mouse neutrophils in the blood at steady state and 24hr after LPS nebulization. (b) Frequencies of human lineages in the blood at steady state and 24hr after LPS nebulization. (c) Numbers of human lineages in the blood at 24hr after LPS nebulization. Data pooled from at least 2 independent experiments. All mice were irradiated with 150 Rads and intra-hepatically injected with 20,000 human fetal liver CD34+ cells at 1-3 days after birth.

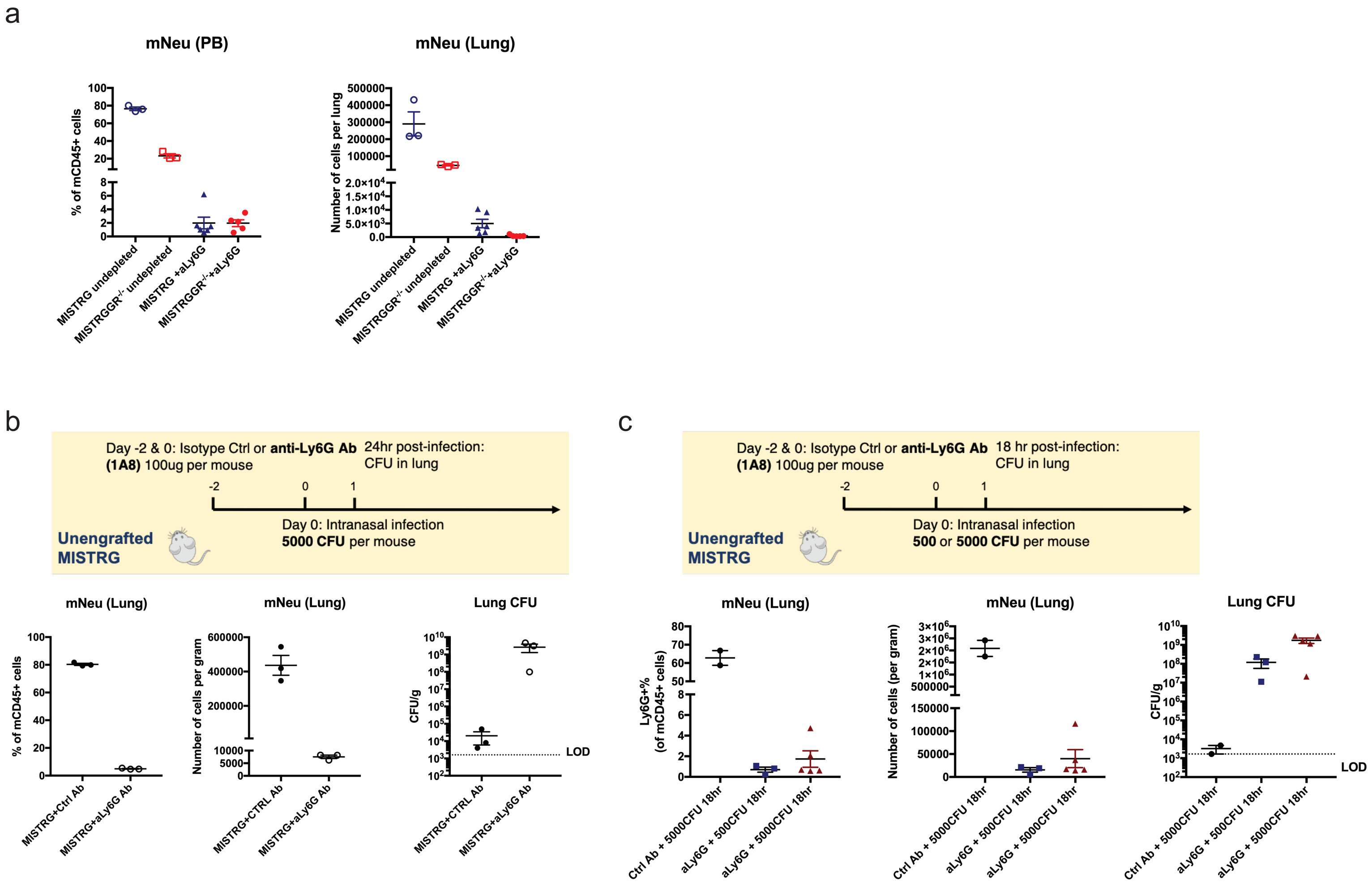


Figure S5. Depletion of murine neutrophils with anti-Ly6G antibody. (a) Validation of murine neutrophil depletion after anti-Ly6G (1A8) antibody treatment. Blood and lung were analyzed at 18 hr post-infection (500 CFU) with untreated engrafted mice as control. Left: Frequencies of mouse neutrophils (mCD45+ Ly6Cmid SSChi) in the blood; Right: Numbers of mouse neutrophils (mCD45+ Ly6Cmid SSChi) in the lung. (b, c) Non-engrafted MISTRG mice were pre-treated with isotype control and anti-Ly6G (1A8) (2 doses of 100 µg, i.v.) for depletion of murine neutrophils, and then intranasally infected with P. aeruginosa. Frequencies and numbers of murine neutrophils and bacterial CFUs in the lung were quantified. (b) Mice were infected with 5000 CFU and analyzed at 24 hr post-infection. (c) Mice were infected with 500 or 5000 CFU of P. aeruginosa and analyzed at 18 hr post-infection.
